# Breaking Boundaries: New locality records and a reassessment of the conservation status of the Khasi Hill Toad (*Bufoides meghalayanus*) endemic to Meghalaya, Northeast India

**DOI:** 10.1101/2024.12.19.629417

**Authors:** Samrat Sengupta, Esau Asakho Chachei, Bitupan Boruah, Anukul Nath, Abhijit Das

## Abstract

*Bufoides meghalayanus* is a Critically Endangered toad, endemic to East Khasi Hills, Meghalaya, Northeast India. This endemic toad was presumed to be restricted to just three locations, all within 1-2 km from one another around the type locality in Meghalaya. This article gives the first account of three previously unknown populations, extending the known geographic range of the species significantly. Additionally, we include the newly obtained molecular sequences of *B. meghalayanus* in a comprehensive molecular phylogeny with a note on its genetic and morphometric variations within these new populations and its congeneric members.

---

India, a megadiverse country with four biodiversity hotspots harbours many narrow endemic amphibian species, whose conservation is severely impeded from the Wallacean shortfall (a knowledge gap in species distribution; Hortal et al. 2015). This is even more acute for the saxicolous group of species inhabiting habitats with significant forest cover and topographic relief. Genus Bufoides (Pillai and Yazdani 1973) under family *Bufonidae* is one such range-restricted genera, endemic to Northeast India (Frost 2024) with just three nominal members (*Bufoides meghalayanus*, East Khasi Hills, Meghalaya; *B. kempi*, Garo Hills, Meghalaya and *B. bhupathyi*, Mizoram; Figure 1). Of these three, *B. meghalayanus* (the type species) is listed as Critically Endangered (CR) under the IUCN criteria B1ab(iii); C2a(ii) with its range restricted to just three locations, all within 1-2 km from the type locality in Cherrapunji (now Sohra), East Khasi Hills, Meghalaya (see IUCN SSC 2022). Furthermore, with fewer than 100 mature individuals forming one subpopulation, its Extent of Occurrence (EOO) is approximately over 18 km^2^ (Deuti et al. 2012). This endangered toad is presumed to be localized in just one precarious pocket of the karst limestone landscape severely impacted by extensive rock-blasting and stone quarrying (Deuti et al. 2012). However, many a times, the delimitated range limits of such narrow endemics can be incomplete, subjective to geographical sampling bias (Narayanan et al. 2022). In this context, we present new occurrence records of *B. meghalayanus* and spatially project them to provide a geographical context of the range of the species, along with genetic and morphometric variations within these populations.

**Figure 1.**
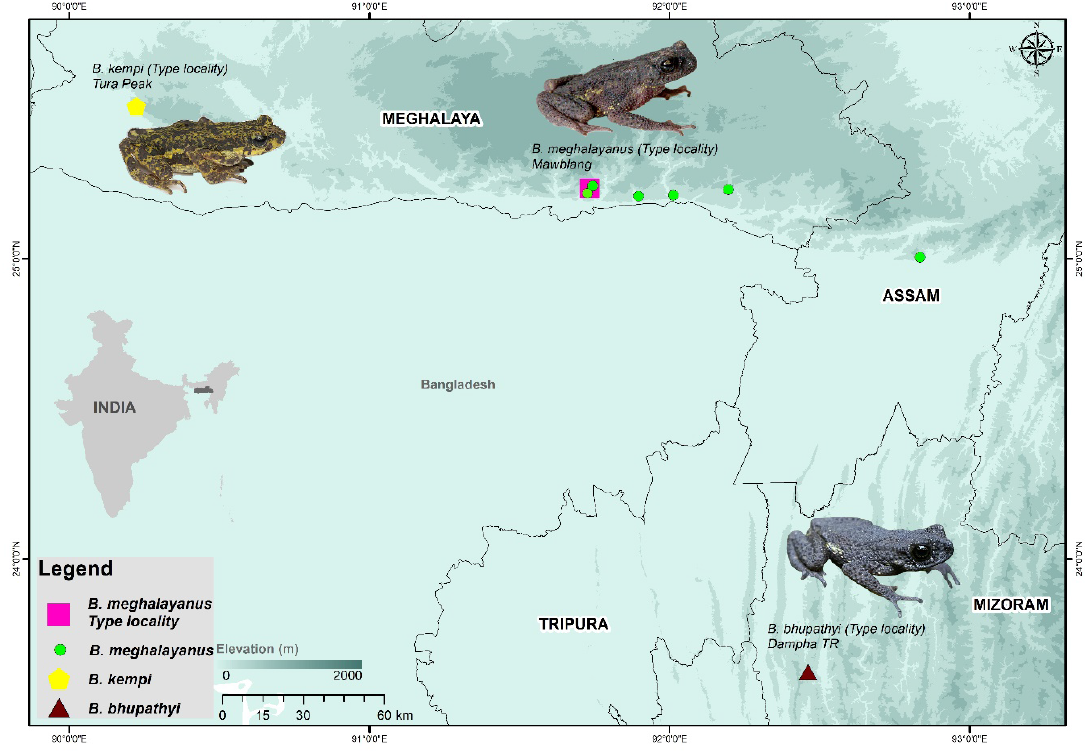
Map of Northeast India showing the distribution of genus *Bufoides* Pillai and Yazdani, 1973, with new locality records for *B. meghalayanus* and the type locality of its three nominal members (*B. meghalayanus*, mawblang, East Khasi Hills, Meghalaya; *B. kempi*, Tura, Garo Hills, Meghalaya and *B. bhupathyi*, Dampa Tiger Reserve, Mizoram).

Building on the findings from previous studies, we broadened our search area incorporating the knowledge gathered from the local community as well as the leading batrachologists working in the landscape. This was followed by an intensive search of potential habitats (see Deuti et al. 2012), accessed with the help of Google Earth imagery. We surveyed across an elevation gradient in Meghalaya (Khasi Hills, Jaintia Hills, and low-lying areas of Ri Bhoi) from April to June 2024. Surveys mostly relied on time-bound “Visual Encounter Surveys” formalized by Crump and Scott (1994), actively searching horizontal slits and rock crevices inside and also within sandstone boulders. A total of 11 potential sites were surveyed over 24 sampling days (0900-2000 hrs) to delineate the potential distribution of the species (Table S1). Representative individuals (Figure 2), when encountered, were collected and the geographic coordinates was marked with a GPS device. Morphometric measurement followed Genomic DNA extraction from the liver samples which were subjected to PCR amplification of a partial fragment of 16S rRNA gene. Metric and meristematic characters (Table 1) were examined by comparing the original description and relevant literature. A principal component analysis (PCA) was performed on the 16 morphometric traits (standardised to their individual SVL) to examine their morphometric distinction from each other. Obtained genetic sequences from the new populations was aligned with 36 other sequences of comparable length belonging to 16 different genera of Asian Toads (Table S2) available at GenBank using MEGA 7.0 software. The resulting alignment was subjected to Maximum Likelihood (ML) phylogenetic reconstruction, with 1000 bootstrap replicates (to evaluate clade stability) under GTR+GAMMA+I model of sequence evolution, using *Limnonectes khasianus* and *Amolops siju* to root the tree (Figure 3).

**Table 1.**
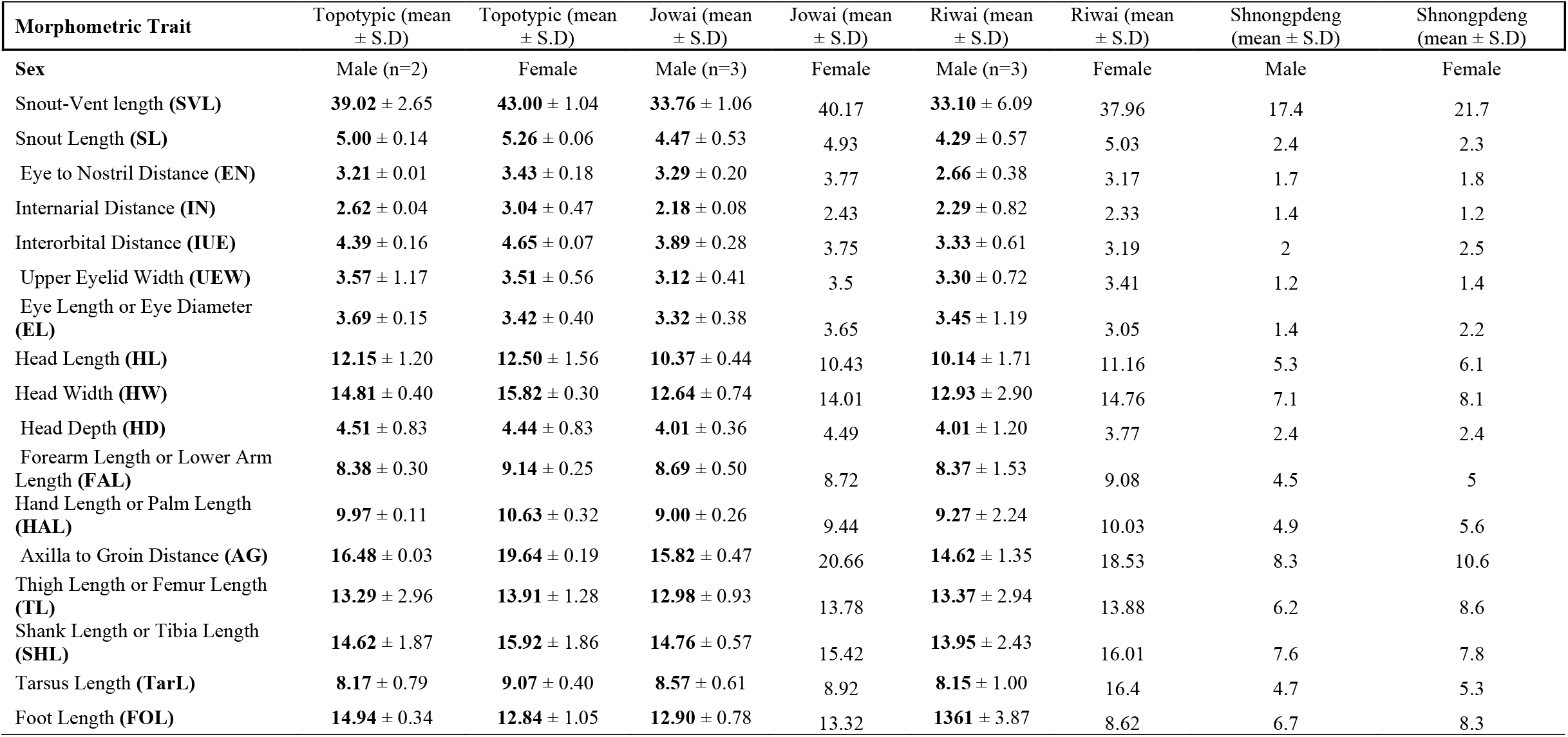
Morphometric details of individuals from the newly recorded populations in the present study.

**Figure 2.**
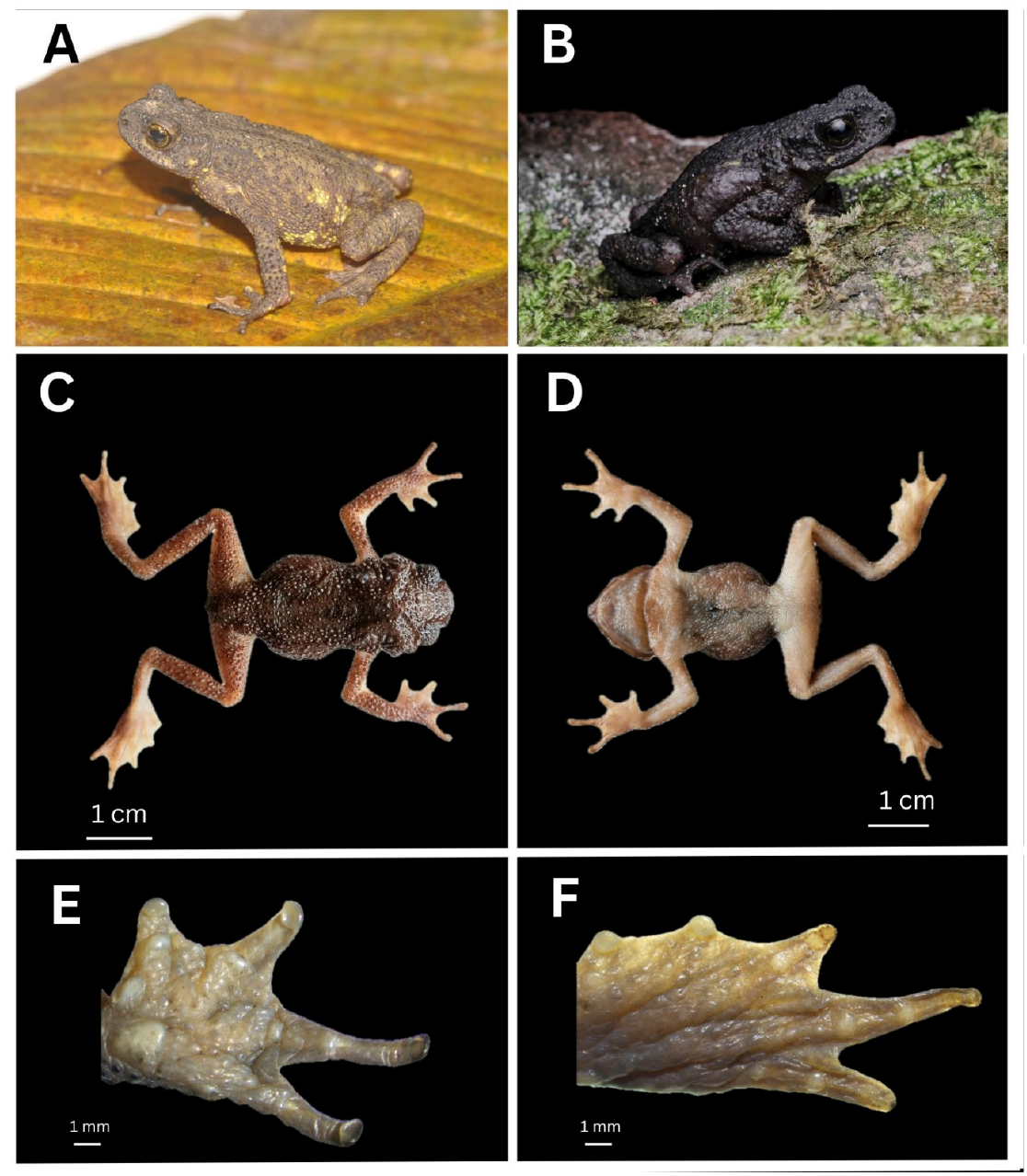
A and B. Male. *B. meghalayanus* in life **C** dorsal view, **D** ventral view, **E** ventral side of the right hand, **F** ventral side of the right feet

**Figure 3.**
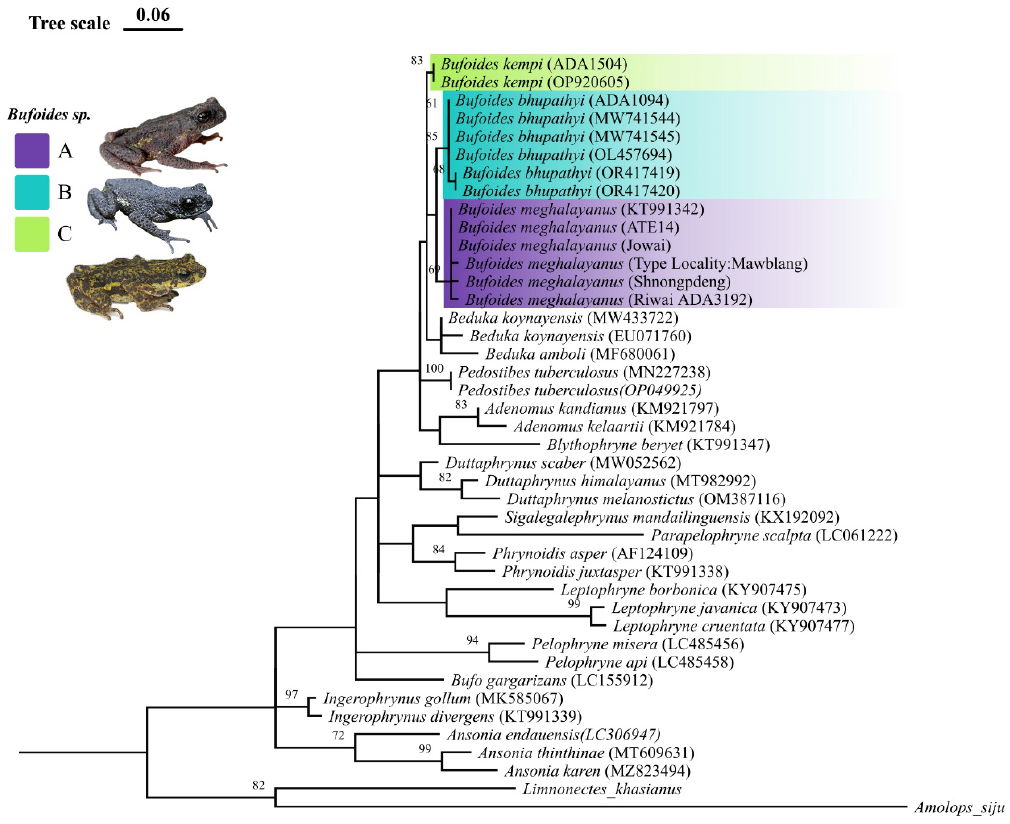
Phylogenetic relationship of the newly recorded populations of *B. meghalayanus* based 16S rRNA.

In total, our survey recorded the presence of three new population of *B. meghalayanus* from Riwai, Shnongpdeng and Jowai (Table S1) which was confirmed by both molecular and morphometric data. Apart from these new records, we also recorded *Bufoides* population from previously documented sites-Mawsmai, Mawblang and Thangkharang area of East Khasi Hills, Meghalaya. All of these sightings were from karst limestone habitat in narrow horizontal rock crevices, pointing to their narrow niche requirement of cool and damp environments. These new populations generally agreed with that of the original as well as that of the expanded description of the species (see Naveen et al. 2022), although the snout-vent length (SVL) of few specimens collected from Riwai turned out to be slightly smaller (∼ 36 mm) than the topotypic specimens (Table 1). Results of the PCA also revealed a significant overlap in the morphometric space among the populations (Figure 4). However, our phylogenetic analysis based on the obtained 16S rRNA partial fragments (∼ 400 to 550 bp in length) from these populations showed minor genetic divergence, despite them clustering into the clade containing the other *Bufoides* species, being nested together with the topotypic material *B. meghalayanus* (Figure 3). The mean genetic distances (p-distance) between the new populations varied by 0.7 – 2.3 % (Table S3), despite their short distance from the type locality (range 17-50 km). Within its congeners, *B. meghalayanus* differs by 3.8 % on 16S (with *B. bhupathyi*) and 6.2 % (with *B. kempi*). Additionally, one opportunistic sighting of a *Bufoides* individual (cf. *B. meghalayanus*) was also recorded from Barail Hill range, Assam (neighbouring state of Meghalaya), and could very well be the first state record of genus *Bufoides* from Assam. These new records extend the easternmost distributional limit of the species ca. 20-30 km (Riwai and Shnongpdeng), extending across the Jaintia Hill Range (Jowai ∼ 50 km) upto the Barail Hill Range (Durbin Tila, Cachar, Assam) from its type locality in Mawblang, East Khasi Hills, Meghalaya. Additionally, these new records slightly lower the elevational limits reported for the species in the country to 100-400 m a.s.l (previously considered to be strictly montane species occurring only on the hilltops between elevations 1,000-1,200 m a.s.l).

**Figure 4.**
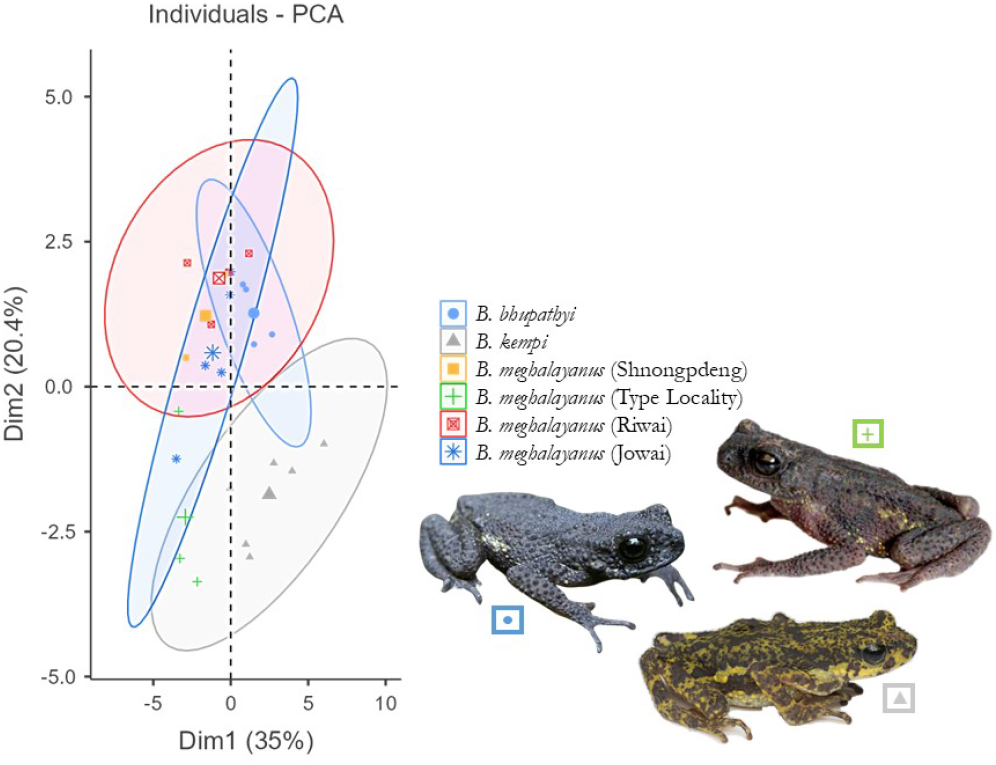
Plot of PCA showing morphometric distinction between the cogeneric members of genus *Bufoides*

Distribution limits for *B. meghalayanus* in Northeast India remain relatively poorly known because they are cryptic and highly seasonal in nature (Pillai and Yazdani 1973). Moreover, only widely scattered sites have been subjected to thorough surveys. Until recently, it was known from only three localities. This is despite the fact that it was first collected and described way back in 1970-71 (named as *Ansonia meghalayana*), only to be re-discovered 30 years later by Das et al. (2009) from around the same locality. However, recent studies including the present one have yielded additional new sites for this critically endangered toad within a span of just three years (2021-2024) and thus illustrate the importance and efficiency of species-focussed surveys when the goal is to delineate the spatial distribution limits of endangered species. Therefore, we urge biologists for more rigorous intensive surveys at a finer scale, combining inputs from ‘Species Distribution model’ and ‘Citizen Science’ to guide future faunal surveys.

The new locality records presented in this study not only extends the known range of the species significantly (Figure 1), but also hint towards a probable broader distribution of the species, and the widely separated locality records might probably reflect inadequate sampling owing to the difficulty of finding these saxicolous toads in Karst landscape of Meghalaya. Albeit this, their specific habitat requirement of narrow horizontal rock crevices in big limestone boulders points to their narrow niche requirement, being widely regarded as a microhabitat specialist (Pillai and Yazdani 1973). This makes them highly susceptible to both anthropogenic habitat alterations and climate change. Also, the newly recorded distribution sites are a major tourist hotspot and the impact of human footprint on potential *Bufoides* habitat also needs to be evaluated as a conservation priority. The Cherrapunji block has seen a concomitant rise in population density from ∼120 people per km^2^ to over 300 individuals per km^2^, between 1961 and 2011 (Census of India, 1961; 2011) adding additional anthropogenic pressure to these relict karst limestone habitats (which are mostly under private land-ownership). This fragile landscape is also undergoing unprecedented rates of deforestation (associated with charcoal production for iron smelting and shifting cultivation) and tourism-induced land use changes along with infrastructural development which has escalated the intensity of limestone mining through extensive stone-blasting and quarrying activities (Prokop 2020). Ideal *Bufoides* habitat around the Mawmluh-Cherra area has been completely destroyed because of such rampant mining activities.

In order to mitigate the detrimental effects posed by stone mining and quarrying, it is crucial to implement stringent regulations and promote sustainable land-use practices. Necessary and proactive measures must be taken to educate local communities and foster community-based conservation alongside periodic population monitoring activities in sites with good stronghold population of *Bufoides*. Evoking a sense of cultural connection as its habitat are in sacred groves (in Khasi ∼ *Law Kyntang*) can strengthen existing community-based conservation model locally.

## Acknowledgments

We extend our gratitude to Meghalaya Forest Department, and the Office of The Additional Principal Chief Conservation of Forests, Wildlife and Chief Wildlife Warden, Meghalaya, Shillong, for permitting us to collect *Bufoides* samples for our research (Permission: **Memo. no. FOR.77/2019/94-A**). We would also like to express our gratitude to our funding agency-Prakriti Fellowship for providing the financial support that enabled us to conduct this research.

## Notes

### Competing Interest Statement

The authors have declared no competing interest.

## References

Census of India, 1991. District Census Handbook, East Khasi Hills District, Village & Town Directory, Parts XI. A&B, Series 16, Director of Census Operations. Meghalaya.

Census of India, 2011. District Census Handbook, East Khasi Hills, Part XII B, Series 18. Directorate of Census Operations, Meghalaya.

Crump ML, Scott NJJ. 1994. Visual encounter surveys. In Heyer WR, Donnelly MA, McDiarmid RW, et al., editors. Measuring and monitoring biological diversity—standard methods for amphibians. Smithsonian Instituition Press. pp.84–92

Das I, Rangad D, Tron RKL, et al. 2009. Rediscovery of the endangered Khasi Hills rock toad, Bufoides meghalayana in Meghalaya, Northeastern India. Froglog 92: 1–4.

Deuti K, Ray S, Dey SK. 2012. Status survey of the Khasi Hills rock toad (Bufoides meghalayana) at Cherrapunjee, Meghalaya. Records of the Zoological Survey of India 92: 21–25.

Frost DR. 2024. Amphibian Species of the World: An Online Reference. Version 6.2. Electronic Database. American Museum of Natural History, New York, USA. http://research.amnh.org/herpetology/amphibia/index.html [Accessed 28 June 2024]

Hortal J, de Bello F, Diniz-Filho JAF, et al. 2015. Seven shortfalls that beset large-scale knowledge of biodiversity. Annual review of ecology, evolution, and systematics 46(1): 523–549

IUCN SSC Amphibian Specialist Group. 2022. Bufoides meghalayanus. The IUCN Red List of Threatened Species 2022.

Narayanan S, Bhat HNP, Paran D, et al. 2022. Truly Absent or Sampling Gaps? Insights on the Potential Distribution of Duttaphrynus hololius (Günther, 1876) from Peninsular India. Current Herpetology 41(2):215–229.

Naveen RS, Chandramouli SR, Kadam G, et al. 2022. Systematics of the enigmatic and narrowly endemic toad genus Bufoides Pillai & Yazdani, 1973: Rediscovery of Bufoides kempi (Boulenger, 1919) and expanded description of Bufoides meghalayanus (Yazdani & Chanda, 1971) (Amphibia: Anura: Bufonidae) with notes on natural history and distribution. Journal of Threatened Taxa 14(12):22277–22292.

Pillai RS, Yazdani GM. 1973. Bufoides, a new genus for the rock-toad, Ansonia meghalayana Yazdani and Chanda, with notes on its ecology and breeding habits. Journal Zoological Society India 25:65–70.

Prokop P. 2020. Remote sensing of severely degraded land: Detection of long-term land-use changes using high-resolution satellite images on the Meghalaya Plateau, northeast India. Remote Sensing Applications: Society and Environment 20:100432.

