## Supplementary Table S1 and S2 for "Breaking Boundaries: New locality records and a reassessment of the conservation status of the Khasi Hill Toad (*Bufoides meghalayanus*) endemic to Meghalaya, Northeast India"

Supplementary Table S1. List of sites surveyed to delineate *Bufoides meghalayanus* distribution limit across Meghalaya, India

| Sl. No | Survey Sites | Elevation (in m) | Effort (in km$^2$) | No. of Individuals | Encounter Rate | Microhabitat |
|---|---|---|---|---|---|---|
| 1 | Mawblang, East Khasi Hills | 1139 | 8 | 11 | 1.4 | Dry Stream |
| 2 | Mawsmai, East Khasi Hills | 1229 | 1 | 0 | - | Dry Stream |
| 3 | Mawsynram, East Khasi Hills | 1071 | 1 | 0 | - | Dry Stream |
| 4 | Thangkharang, East Khasi Hills | 862 | 6 | 13 | 2.2 | Forest Patch |
| 5 | Khliehriat, East Jaintia Hills | 717 | 1 | 0 | - | Dry Stream |
| 6 | Kudengrim, West Jaintia Hills | 686 | 0 | 0 | - | Forest Patch |
| 7 | Umden, Ri Bhoi | 540 | 1 | 0 | - | Forest Patch |
| 8 | Jowai, West Jaintia Hills | 654 | 5 | 17 | 3.4 | Dry Stream |
| 9 | Sohbar, East Khasi Hills | 513 | 1 | 0 | - | Dry Stream |
| 10 | Riwai, East Khasi Hills | 425 | 5 | 4 | 0.8 | Dry Stream |
| 11 | Shnongpdeng, Dawki, West Jaintia Hills | 149 | 5 | 2 | 0.4 | Dry Stream |

Supplementary Table S2. List of sequences used in the molecular analyses in this study with their GenBank accession numbers and localities.

| Sl. No | Taxon | Range/ Collection Location | NCBI No. | Reference |
|---|---|---|---|---|
| 1 | *Adenomus kandianus* | Endemic to Sri Lanka | KM921797 | Meegaskumbura et al. 2016 |
| 2 | *Adenomus kelaartii* | Endemic to Sri Lanka | KM921784 | Meegaskumbura et al. 2015 |
| 3 | *Ansonia endauensis* | Endemic to the Malay Peninsula | LC306947 | Unpublished |
| 4 | *Ansonia karen* | Western Thailand | MZ823494 | Suwannapoom et al. 2021. |
| 5 | *Ansonia thinthinae* | Endemic to Myanmar | MT609631 | Unpublished |
| 6 | *Beduka amboli* | Endemic to Western Ghats, India | MF680061 | Unpublished |
| 7 | *Beduka koynayensis* | Endemic to Western Ghats, India | EU071760 | Unpublished |
| 8 | *Beduka koynayensis* | | MW433722 | Dinesh et al. 2023 |
| 9 | *Blythophryne beryet* | India (Andaman and Nicobar Islands) | KT991347 | Chandramouli et al. 2016 |
| 10 | *Bufo gargarizans* | Endemic to East Asia | LC155912 | Unpublished |
| 11 | *Bufoides bhupathyi* | Endemic to Northeast India | OR417419 | Naveen et al. 2023 |
| 12 | *Bufoides bhupathyi* | | OR417420 | Naveen et al. 2024 |
| 13 | *Bufoides bhupathyi* | | ADA1094 | Unpublished |
| 14 | *Bufoides bhupathyi* | | MW741544 | Unpublished |
| 15 | *Bufoides bhupathyi* | | MW741545 | Unpublished |
| 16 | *Bufoides bhupathyi* | | OL457694 | Unpublished |
| 17 | *Bufoides kempi* | Endemic to Northeast India | ADA1504 | Unpublished |
| 18 | *Bufoides kempi* | | OP920605 | Naveen et al. 2022 |
| 19 | *Bufoides meghalayanus* | Endemic to Northeast India | ADA3192 | Unpublished |
| 20 | *Bufoides meghalayanus* | | KT991342 | Chandramouli et al. 2016 |
| 21 | *Bufoides meghalayanus* | | **Shnongpdeng** | Unpublished |
| 22 | *Bufoides meghalayanus* | | **Type Locality** | Unpublished |
| 23 | *Bufoides meghalayanus* | | **Jowai** | Unpublished |
| 24 | *Bufoides meghalayanus* | | PQ305648/ATE14 | Unpublished |

| | | | | |
|---|---|---|---|---|
| 25 | *Duttaphrynus himalayanus* | Widely Spread throughout Himalayan mountains | MT982992 | Khatiwada et al. 2021 |
| 26 | *Duttaphrynus melanostictus* | South and Southeast Asia | OM387116 | Unpublished |
| 27 | *Duttaphrynus scaber* | India and Sri Lanka | MW052562 | Ashaharraza et al. 2020 |
| 28 | *Ingerophrynus divergens* | Brunei, Indonesia, Malaysia, Thailand | KT991339 | Chandramouli et al. 2016 |
| 29 | *Ingerophrynus gollum* | Endemic to the Peninsular Malaysia | MK585067 | Kin Onn Chan and L. Lee Grismer (2019) |
| 30 | *Leptophryne borbonica* | Brunei, Indonesia, Malaysia, Thailand | KY907475 | Hamidy et al. 2018 |
| 31 | *Leptophryne cruentata* | Endemic to Java, Indonesia | KY907477 | Hamidy et al. 2018 |
| 32 | *Leptophryne javanica* | Endemic to Indonesia | KY907473 | Hamidy et al. 2018 |
| 33 | *Parapelophryne scalpta* | Endemic to Hainan, China | LC061222 | Matsui et al. 2015 |
| 34 | *Pedostibes tuberculosus* | Endemic to Western Ghats, India | MN227238 | Unpublished |
| 35 | *Pedostibes tuberculosus* | | OP049925 | Unpublished |
| 36 | *Pelophryne api* | Endemic to Indonesia | LC485458 | Eto,K. and Matsui,M 2019 |
| 37 | *Pelophryne misera* | Endemic to Indonesia | LC485456 | Eto,K. and Matsui,M 2019 |
| 38 | *Phrynoidis asper* | Mainland Southeast Asia and the Greater Sundas | AF124109 | Vences et al. 2000 |
| 39 | *Phrynoidis juxtasper* | Borneo (Brunei, Indonesia, and Malaysia) and Sumatra (Indonesia) | KT991338 | Chandramouli et al. 2016 |
| 40 | *Sigalegalephrynus mandailinguensis* | Endemic to Indonesia | KX192092 | Smart 2017 |
| 41 | *Amolops siju* | OUTGROUP | | |
| 42 | *Limnonectes khasianus* | | | |
